# Protease-activated receptor 1-mediated matrix metalloprotease signalling in sensory neurons

**DOI:** 10.1101/2025.08.20.671259

**Authors:** Katie J Williams, Giulia Galimberti, James P Higham, Emily Overington, Charity N Bhebhe, Tim Raine, Paola Sacerdote, David C Bulmer

**Affiliations:** Department of Pharmacology, University of Cambridge, Cambridge, United Kingdom; Department of Gastroenterology, Addenbrookes Hospital, Cambridge University Teaching Hospitals, Cambridge, United Kingdom; Department of Pharmacological and Biomolecular Sciences, University of Milan, Milan, Italy

## Abstract

Visceral pain is a prevalent and debilitating symptom of inflammatory bowel disease (IBD). However, current pain therapies are often ineffective, raising the possibility that novel disease mediators might be contributing to pain during inflammation. Our study provides new insights into how matrix metalloproteinases (MMPs), which are elevated in IBD, stimulate sensory neurons. We demonstrate that MMP3, MMP8, and MMP9 induce intracellular Ca^2+^ release in dorsal root ganglion (DRG) neurons through activation of protease-activated receptor 1 (PAR1) and subsequent activation of phospholipase C (PLC). Characterisation of the neuronal populations stimulated by these MMPs suggests that a subset is likely nociceptive. In contrast, MMP2 and MMP13, although capable of cleaving PAR1 in other cell types, do not induce Ca^2+^ mobilisation in DRG neurons. Interestingly, pre-treatment with MMP2 or MMP13 reduces the neuronal response to MMP3 or PAR1 agonist, suggesting that MMP2 and MMP13 act on PAR1 in a manner which prevents further activation. Additionally, MMP10 induces Ca^2+^ mobilisation in DRG neurons but through a PAR1-independent mechanism. These findings uncover a previously unrecognised role for MMP signalling in sensory neurons, highlighting a potential mechanism by which MMPs could contribute to the pro-nociceptive environment in the inflamed bowel.

## Introduction

A greater understanding of the mediators and mechanisms underpinning colonic nociception during inflammation will shed light on potential targets for future analgesics for use in inflammatory bowel disease (IBD). Abdominal pain is a distressing and poorly-treated symptom of IBD; current analgesics often lack clinical efficacy and have intolerable side-effects. As such, there is a significant unmet clinical need for new therapies for the treatment of abdominal pain.

Matrix metalloproteases (MMPs) are zinc-dependent endopeptidases which maintain the extracellular matrix, as well as cleaving a diverse set of substrates, including cell surface receptors, cytokines and growth factors. Consequently, MMPs play an important role in tissue repair and remodelling, inflammation and cell migration (Cabral-Pacheco et al., 2020; Nagase et al., 2006). The expression and activity of various members of the MMP family are elevated in the inflamed bowel (Higham et al., 2024; Pedersen et al., 2009), raising the possibility that MMPs contribute to the pathophysiology of IBD (Ravi et al., 2007).

Protease-activated receptor 1 (PAR1) is a G-protein-coupled receptor activated by the endopeptidase-mediated cleavage of its extracellular N-terminal domain, leading to the exposure of a tethered ligand. PAR1 is primarily involved in platelet activation (Andersen et al., 1999) and haemostasis (Nieman, 2016), but has also been shown to contribute to pro-nociceptive signalling (Desormeaux et al., 2018) and IBD pathophysiology (Saeed et al., 2017). Furthermore, some members of the MMP family, such as MMP1, can cleave and activate PAR1, representing a potentially novel pro-nociceptive signalling pathway in the inflamed bowel.

In this study, we used Ca^2+^ imaging of sensory neuronal soma to probe the effects of MMPs which exhibit elevated expression in colonic biopsies from patients with Crohn’s disease. We found that MMP3, 8 and 9 evoke a PAR1-dependent increase in intracellular Ca^2+^ concentration in a population of small-diameter sensory neurons. Conversely, MMP10 raised intracellular Ca^2+^ concentration independent of PAR1. Finally, MMP2 and 13 did not raise intracellular Ca^2+^ concentration in sensory neurons, though pre-treatment with either MMP2 or 13 inhibited the subsequent response to MMP3 or TRAP6 (a synthetic PAR1 agonist). These data demonstrate that MMPs may contribute to pro-nociceptive signalling in the inflamed bowel, and that PAR1 plays a significant role in MMP signalling in sensory neurons.

## Methods

### Preparation of sensory neurons

Sensory neurons were isolated from the dorsal root ganglia (DRG) from male C57Bl/6 mice aged between 8-14 weeks. All animal work was carried out in accordance with Schedule 1 of the Animals (Scientific Procedures) Act 1986 Amendment Regulations 2012. Mice were housed in groups of up to six littermates under a 12-hour light-to-dark cycle with *ad libitum* access to food, water, bedding and enrichment (e.g., tunnels, chews).

Neurons were cultured as described previously (Barker et al., 2022; Higham et al., 2024). DRG were harvested from spinal levels T12-L6. After removal, DRG were incubated (37°C, 5% CO_2_) first with 1 mg/mL type 1A collagenase (Sigma Aldrich, St Louis, MA, USA) for 15 minutes and second with 1 mg/mL trypsin (Sigma Aldrich) for 30 minutes, both of which were prepared in Lebovitz L-15 GlutaMAX media (Invitrogen, Waltham, MA, USA) supplemented with 6 mg/mL bovine serum albumin (Sigma Aldrich) and 2.6% vol/vol NaHCO_3_ (Invitrogen). Following enzymatic digestion, DRG were triturated with a P1000 pipette tip and centrifuged at 100*g* for 30 seconds. Trituration and centrifugation were repeated five times, with the supernatant (containing dispersed neurons) collected each time. The collected supernatant was centrifuged (100*g*) a final time for 5 minutes, and the pelleted cells were resuspended in supplemented L-15 media (10% vol/vol foetal bovine serum, 2.6% vol/vol NaHCO_3_, 1.5% vol/vol D-glucose and 2% vol/vol penicillin/streptomycin; all purchased from Invitrogen). Dispersed neurons were plated onto glass-bottomed dishes (MatTek, Ashland, MA, USA) coated with laminin and poly-D-lysine. Neurons were incubated at 37°C in 5% CO_2_ and used for imaging 48 hours after plating.

### Ca^2+^ imaging

To load neurons with Ca^2+^ indicator (Fluo-4), they were incubated with Fluo-4-AM (10 μM; ThermoFisher Scientific, Waltham, MA, USA) diluted in extracellular bath solution (in mM: 140 NaCl, 4 KCl, 1 MgCl_2_, 2 CaCl_2_, 4 D-glucose and 10 HEPES; pH adjusted to 7.35-7.45 with NaOH) for 40 minutes (room temperature; shielded from light). Following this, neurons were washed with, and bathed in, extracellular solution, and dishes were mounted on the stage of an inverted microscope (Nikon Eclipse TE2000S). For experiments in which the effect of an antagonist was investigated, neurons were preincubated with antagonist-containing solution for 10 minutes prior to imaging (antagonists were also present throughout imaging). To identify non-peptidergic neurons in culture, neurons were incubated with isolectin-B4 (IB4, conjugated to Alexa Fluor 568; 10 μg/mL; ThermoFisher Scientific) for 10 minutes prior to imaging (with an additional rinse to wash off unbound IB4).

Fluo-4 was excited using a 470 nm LED (Cairn Research, Kent, UK). Emission at 520 nm was captured using a CCD camera (Retiga Electro, Photometrics, Tucson, AZ, USA) at 2.5 fps with 100 ms exposure and recorded using μManager. During imaging, neurons were superfused with extracellular bath solution at ∼0.5 mL/minute using a gravity-fed, valve-controlled perfusion system. Where multiple drugs were applied to the same dish, a 4-minute wash was applied between applications. At the end of each experiment, 50 mM KCl was applied to identify viable neurons and allow the normalisation of recorded fluorescence values.

Using ImageJ (v2; NIH, MA, USA), regions of interest (ROI) were manually circled around individual neurons from a brightfield image and overlaid onto fluorescent images. Pixel intensity was measured and analysed with custom-written scripts (Barker et al., 2022; Higham et al., 2024) in RStudio (RStudio, MA, USA) to calculate the proportion of neurons stimulated by each drug application and the corresponding magnitude of response. Background fluorescence was subtracted from each region of interest, and fluorescence intensity (F) was baseline-corrected (accounting for varying baselines between neurons) and normalised to the peak fluorescence value during KCl application (F/F_pos_). Cells were excluded from analysis if their fluorescence did not increase by >5% over baseline during KCl application. Neurons were classed as responsive to a particular drug if fluorescence >0.1 F/F_pos_ was reached during drug application.

### Statistics

Data were scrutinised to ensure they met the assumptions of parametric statistical tests. Normality was assessed using the Shapiro-Wilk test and homogeneity of variances was assessed using the Brown-Forsythe test. Where the assumptions of parametric tests were not met, non-parametric, rank-based alternatives were used. Details of sample sizes and statistical tests are given in figure legends. Statistical analyses were carried out in Prism (v10.2.0; GraphPad Inc, La Jolla, CA, USA). Reporting of data in this study is in accordance with the ARRIVE guidelines.

## Results

### Matrix metalloprotease expression is elevated in Crohn’s disease

We used our previously-published bulk RNAseq analysis of human colonic biopsies (Higham et al., 2024) to probe the potential role of MMPs in the inflamed bowel. Multiple members of the MMP family of proteases were found to be upregulated in colonic biopsies from patients with Crohn’s disease. In particular, the expression of MMP1, 2, 3, 8, 9, 10 and 13 was found to be elevated compared to non-inflamed control biopsies (Figure 1A). In line with this, enrichment analysis (GO Molecular Function) of the set of genes upregulated in Crohn’s disease highlighted the enrichment of multiple gene ontology terms relating to peptidase activity, including metallopeptidase activity (Figure 1B). Other enriched gene ontology terms included cytokine and chemokine receptor activity, tyrosine kinase activity, and vascular endothelial growth factor receptor activity.

**Figure 1:**
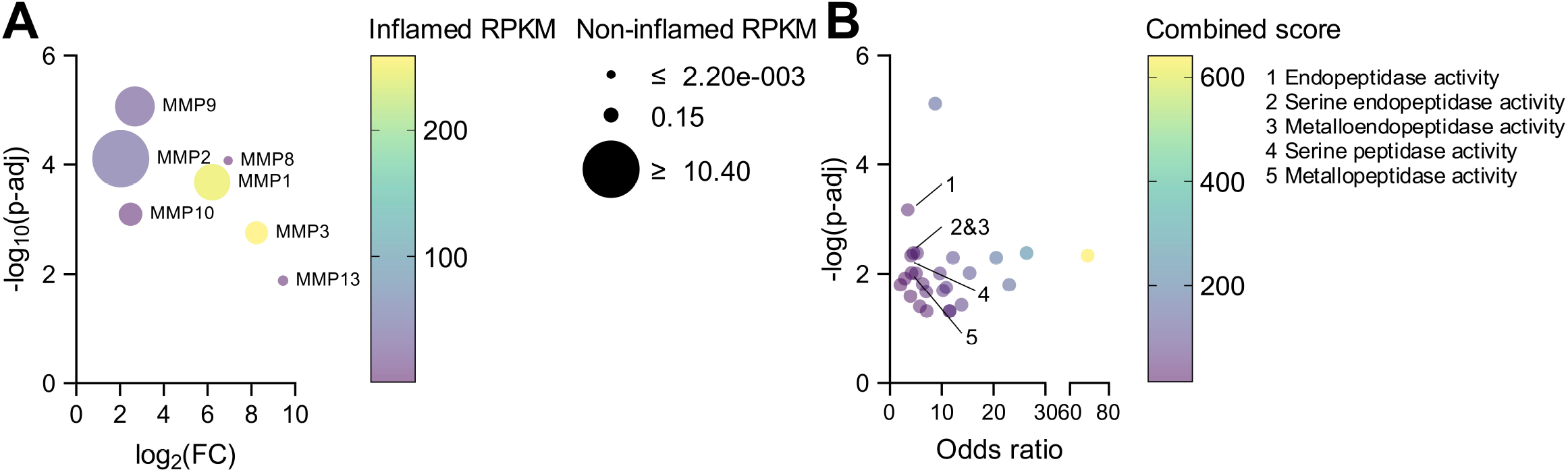
Elevated expression of MMPs in Crohn’s disease (data redrawn from Higham *et al*., 2024). (A) Bubble plot showing the fold-change in expression (expressed as log_2_ compared to non-inflamed controls) and the corresponding p-value (compared to non-inflamed controls, Benjamini-Hochberg corrected) for MMP1, 2, 3, 8, 9, 10 and 13. The size of the bubbles indicates the expression (reads per kilobase per million, RPKM) of each MMP in non-inflamed biopsies, and bubble colour indicates expression in Crohn’s disease biopsies. (B) Bubble plot showing the enrichment of gene ontology terms related to molecular function (annotated gene set: GO Molecular Function), with those specifically related to protease activity highlighted. The odds ratio is the odds of finding a gene annotated to a particular molecular function in the upregulated gene set compared to the odds of finding a gene annotated to a particular molecular function in the background. Significance of gene ontology term enrichment was determined using Fisher’s exact test with Benjamini-Hochberg correction.

The expression of several other MMPs was also raised in Crohn’s disease (MMP7, 12, 14, 16 and 25), but these were not selected for investigation in the present study. These data are in agreement with previous work documenting the elevated expression of MMPs in the inflamed bowel (Von Lampe et al., 2000).

### Matrix metalloproteases 3, 8 and 9 raise cytosolic [Ca^2+^] in sensory neurons

To begin to characterise MMP-evoked Ca^2+^ signals, we applied increasing concentrations of MMPs to Fluo4-loaded sensory neurons. We found that MMP3, 8 and 9 evoked Ca^2+^ signals in a subset of sensory neurons. The application of 0.1 nM of MMP3, 8 or 9 stimulated a proportion of neurons equivalent to that of vehicle (Figure 2A). A greater proportion of neurons was stimulated by 1 nM of each MMP: 19.6±2.5% (F(3, 16) = 6.27, p = 0.0086 compared to vehicle, Figure 2A *left*), 25.5±3.2% (F(3, 16) = 20.48, p < 0.0001 compared to vehicle, Figure 2A *middle*) and 16.5±1.1% (F(3, 16) = 9.27, p = 0.0053 compared to vehicle, Figure 2A *right*) of neurons were stimulated by MMP3, 8 and 9, respectively. Given that the application of 10 nM of each MMP yielded no further increase in response (Figure 2A), 1 nM was used for all subsequent experiments.

**Figure 2:**
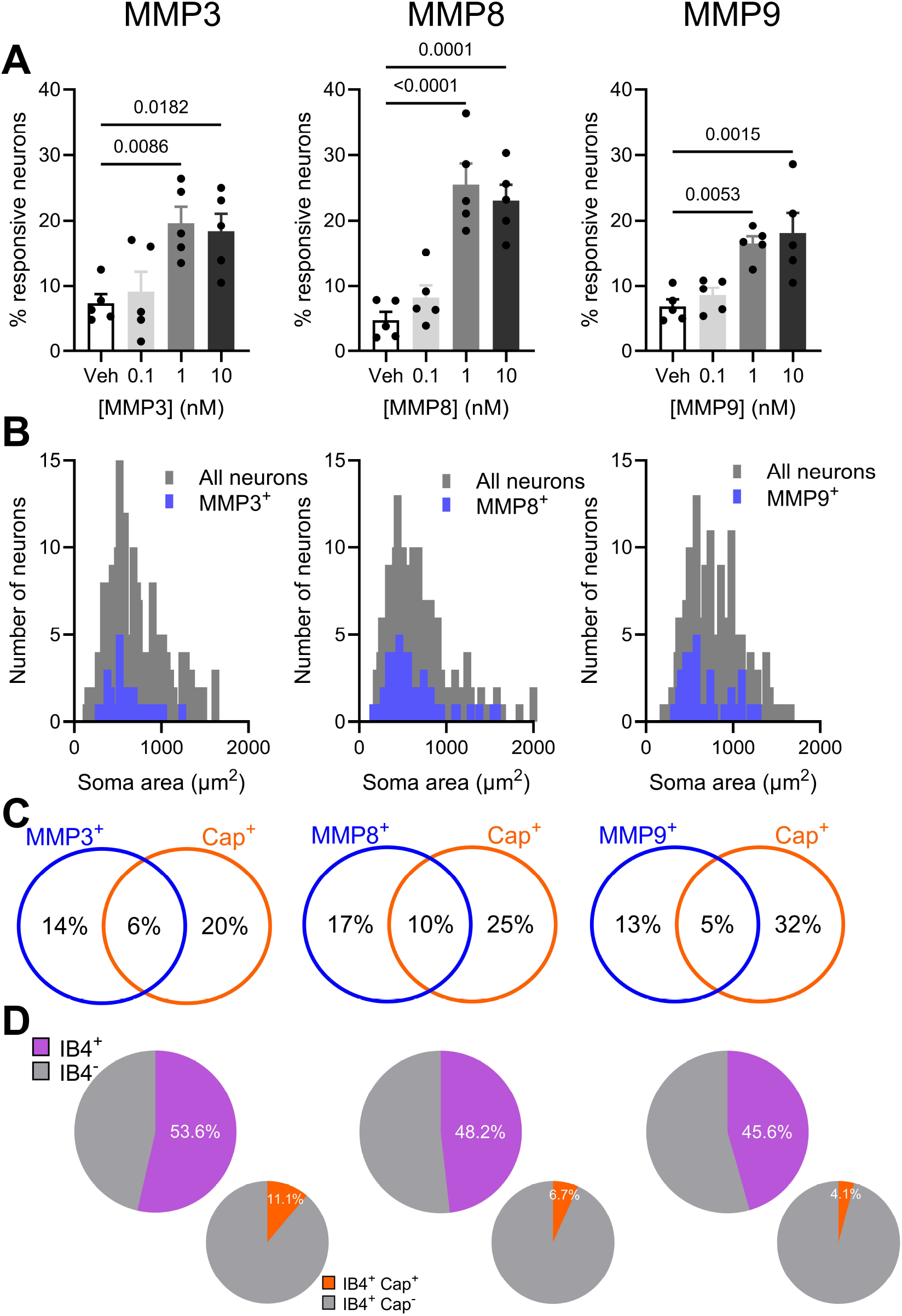
Properties of MMP-induced Ca^2+^ signals in sensory neurons *in vitro*. (A) MMP3 (left), MMP8 (middle) and MMP9 (right) stimulate sensory neurons in a concentration-dependent manner (5 independent experiments per group, 21-78 neurons per experiment). One-way ANOVA with Dunnett’s *post-hoc* tests. (B) Soma size histograms showing the area of sensory neurons sensitive to MMP3 (left), MMP8 (middle) and MMP9 (right). MMP3, 222 neurons; MMP8, 266 neurons; MMP9, 219 neurons. (C) Venn diagrams (not to scale) showing the co-sensitivity between MMP3 (left), MMP8 (middle) and MMP9 (right) and capsaicin. (D) (*Top*) Charts showing the proportion of sensory neurons sensitive to MMP3 (left), MMP8 (middle) and MMP9 (right) which were positively stained by the non-peptidergic neuronal marker, IB4. (*Bottom*) Charts showing the proportion of IB4-positive, MMP-sensitive neurons which are also co-sensitive to capsaicin. 3 independent experiments, 20-65 neurons per experiment.

We probed the identity of MMP-responsive sensory neurons by examining their soma size and co-sensitivity to capsaicin, an agonist of TRPV1 (a key marker of a subset of nociceptive neurons). The vast majority (∼90%) of the neurons sensitive to MMP3, 8 and 9 exhibited a soma area less than 1000 μm^2^ (Figure 2B). MMP3-, 8- and 9-sensitive neurons exhibited a median soma area of 536 μm^2^ (interquartile range, IQR: 421-708 μm^2^), 497 μm^2^ (IQR: 408-679 μm^2^) and 581 μm^2^ (IQR: 487-816 μm^2^), respectively. The sequential application of MMP and capsaicin (1 μM) revealed three neuronal populations: those which responded to only MMP, those which responded to only capsaicin, and those which responded to both MMP and capsaicin. Approximately one-third of the neurons responsive to each MMP were co-sensitive to capsaicin (Figure 2C). We also used staining with the non-peptidergic neuronal marker, IB4, to further characterise MMP-sensitive neurons. In total, of 331 sensory neurons imaged, 136 (41.1%) were positively-stained with IB4. Within each of the MMP-sensitive populations, around one-half were IB4-positive, with the majority (∼90%) of these IB4-positive neurons insensitive to capsaicin (Figure 2D). It is important to note that we did not observe any change in the neuronal sensitivity to capsaicin following pre-treatment with MMP3, 8 or 9. That said, due to the high concentration of capsaicin used (1 μM), these experiments were not optimised to detect increases in the response to capsaicin.

### Matrix metalloprotease signalling via protease-activated receptor 1

To investigate the potential role of PAR1 in MMP-evoked Ca^2+^ signals, we first verified functional expression of PAR1 in sensory neurons in culture and validated the effect of a PAR1 antagonist. Application of TRAP6 (1 μM) evoked a rise in intracellular [Ca^2+^] in 28.6±5.7% of sensory neurons, compared to 3.6±1.4% following the application of vehicle alone (F(2, 12) = 14.53, p = 0.0008, Figure 3). The proportion of TRAP6-sensitive neurons was reduced by pre-incubation with vorapaxar (1 μM), a PAR1-selective antagonist. In the presence of vorapaxar, TRAP6 evoked a rise in intracellular [Ca^2+^] in only 8.5±1.2% of sensory neurons (p = 0.0045 compared to TRAP6 alone, p > 0.99 compared to vehicle, Figure 3).

**Figure 3:**
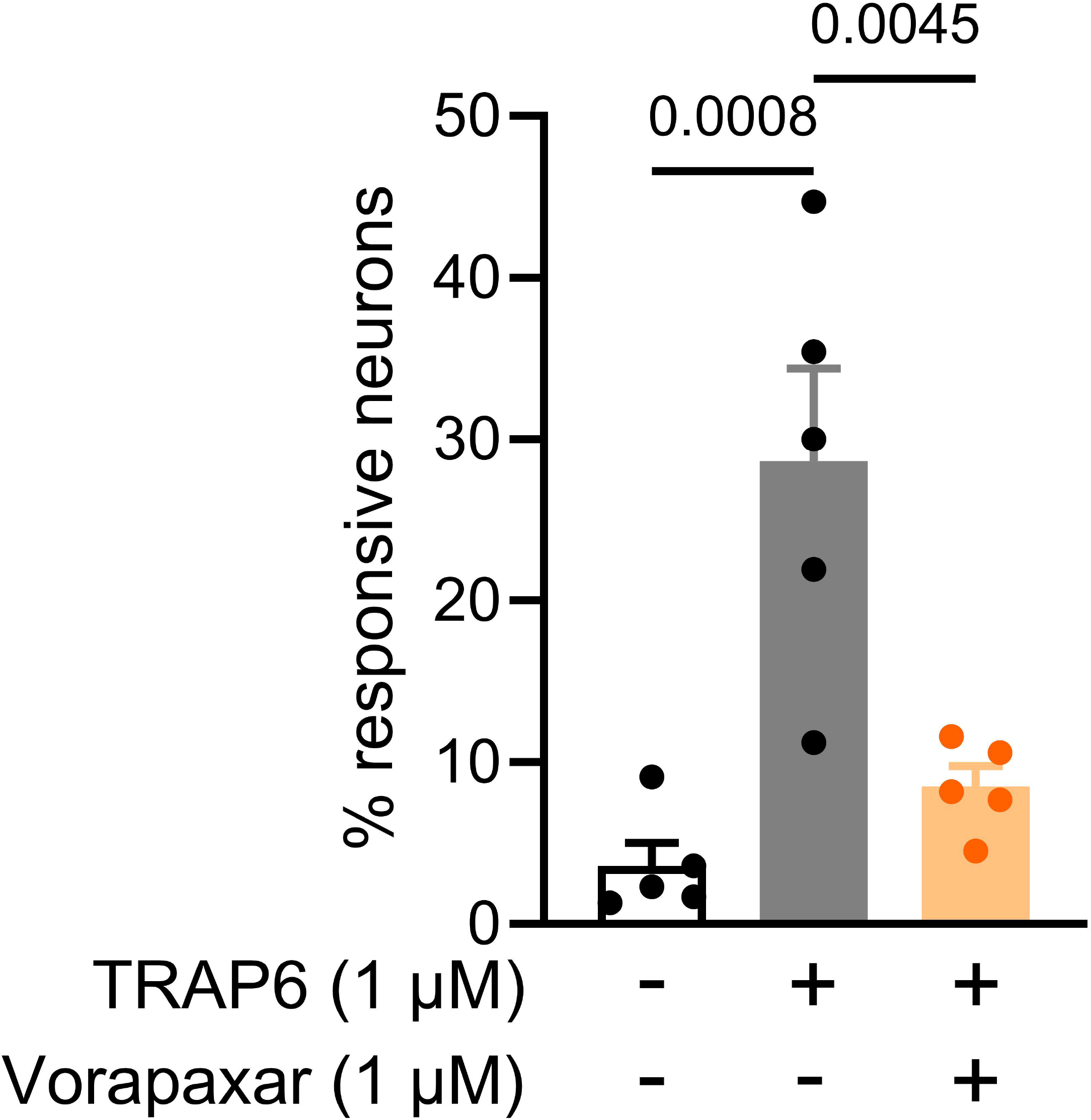
Functional expression of PAR1 in sensory neurons. Grouped data showing the proportion of neurons stimulated by vehicle, TRAP6, and TRAP6 after treatment with vorapaxar (5 independent experiments per group, 21-78 neurons per experiment). One-way ANOVA with Bonferroni’s *post-hoc* tests.

We next tested whether vorapaxar inhibited the responses elicited by MMP3, 8 and 9 (1 nM). As in our previous experiments (see Figure 2), MMP3, 8 and 9 stimulated a rise in intracellular [Ca^2+^] in 16.5±1.0%, 21.1±1.6% and 16.5±1.1% of sensory neurons, respectively (Figure 4A). The proportion of MMP3-, 8- and 9-responsive neurons was reduced to 5.2±1.5% (F(2, 12) = 34.30, p < 0.0001), 11.4±2.1% (F(2, 12) = 23.53, p = 0.0046) and 7.1±0.6% (F(2, 12) = 44.37, p < 0.0001), respectively, by pre-treatment with vorapaxar (Figure 4A). These results indicate that MMP3, 8 and 9 cleave PAR1. We also found that MMP1 stimulated a subset of sensory neurons (13.0±2.3%), but a higher concentration (10 nM) was required. The response to MMP1 was inhibited by another PAR1 antagonist, SCH79797 (10 μM, p = 0.024), but MMP1 was not investigated further. In contrast to MMP1, 3, 8 and 9, the application of MMP10 (1 nM) stimulated a rise in intracellular [Ca^2+^] in 18.7±1.9% of sensory neurons (F(2,12) = 17.12, p = 0.0005 compared to vehicle), but this was not inhibited by pre-treatment with vorapaxar (p > 0.99 compared to MMP10 alone, p = 0.0017 compared to vehicle, Figure 5).

**Figure 4:**
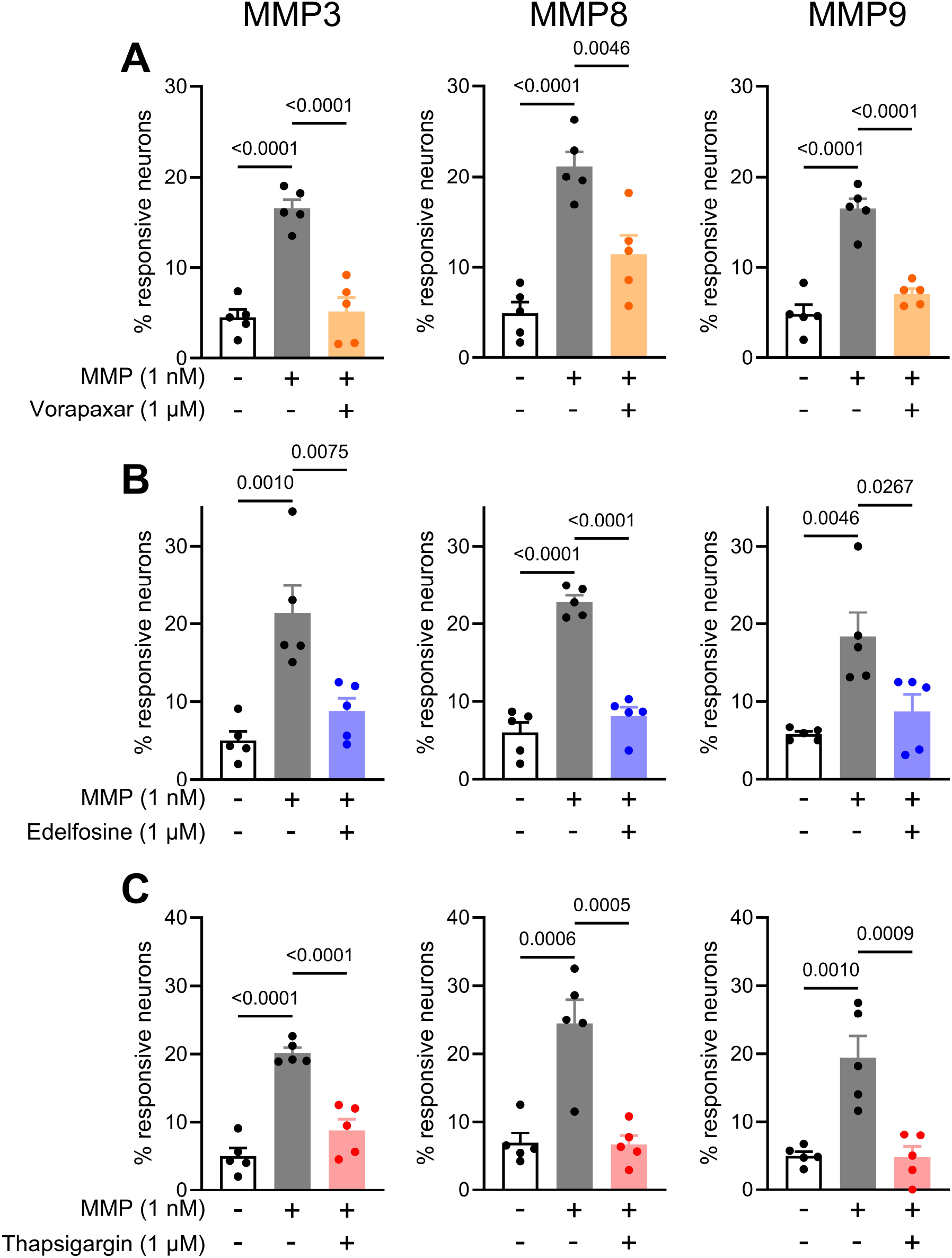
MMP3-, 8- and 9-induced Ca^2+^ signals depend on PAR1 and PLC-mediated Ca^2+^ release from intracellular stores. (A) Grouped data showing the proportion of neurons stimulated by vehicle, MMP alone MMP3, *left*; MMP8, *middle*; MMP9, *right*), and MMP after treatment with vorapaxar (5 independent experiments per group, 20-76 neurons per experiment). One-way ANOVA with Bonferroni’s *post-hoc* tests. (B) Grouped data showing the proportion of neurons stimulated by vehicle, MMP alone MMP3, *left*; MMP8, *middle*; MMP9, *right*), and MMP after treatment with edelfosine (5 independent experiments per group, 21-64 neurons per experiment). One-way ANOVA with Bonferroni’s *post-hoc* tests. (C) Grouped data showing the proportion of neurons stimulated by vehicle, MMP alone MMP3, *left*; MMP8, *middle*; MMP9, *right*), and MMP after treatment with thapsigargin (5 independent experiments per group, 23-72 neurons per experiment). One-way ANOVA with Bonferroni’s *post-hoc* tests.

**Figure 5:**
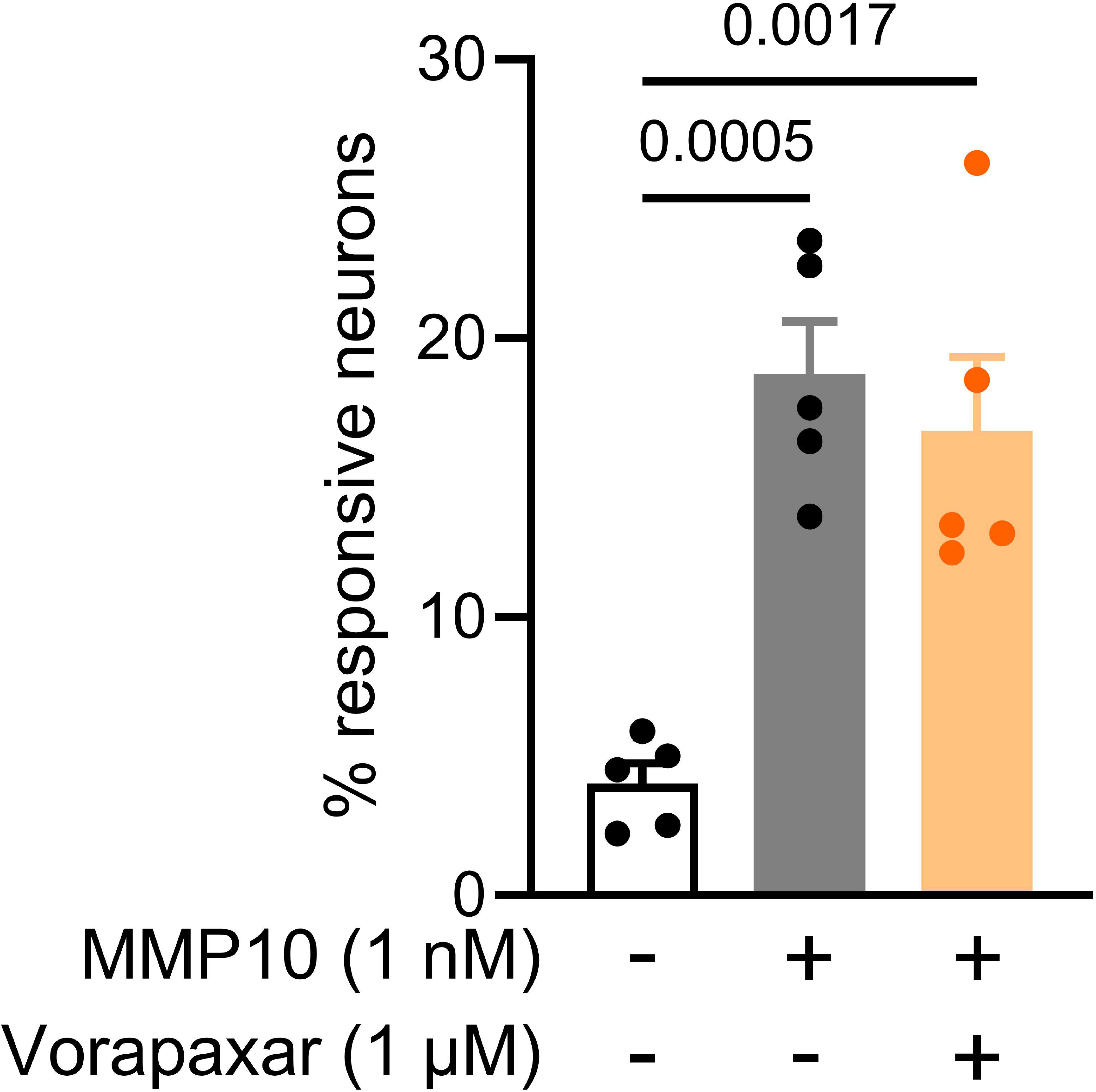
MMP10-evoked Ca^2+^ signals do not depend on PAR1. Grouped data showing the proportion of neurons stimulated by vehicle, MMP10, and MMP10 after treatment with vorapaxar (5 independent experiments per group, 22-50 neurons per experiment). One-way ANOVA with Bonferroni’s *post-hoc* tests.

To further probe the intracellular signalling evoked by MMP3, 8 and 9, we used edelfosine (1 μM) and thapsigargin (1 μM) to test whether Ca^2+^ signals were dependent on phospholipase C (PLC) and Ca^2+^ release from intracellular stores. Pre-treatment with edelfosine reduced the proportion of neurons responding to application of MMP3 (F(2, 12) = 13.48, p = 0.0075), MMP8 (F(2, 12) = 66.05, p < 0.0001) and MMP9 (F(2, 12) = 9.08, p = 0.027, Figure 4B). Similarly, depletion of intracellular Ca^2+^ stores by pre-treatment with thapsigargin also attenuated the proportion of neurons responding to MMP3 (F(2, 12) = 40.80, p < 0.0001), MMP8 (F(2, 12) = 19.10, p = 0.0005) and MMP9 (F(2, 12) = 16.63, p = 0.0009, Figure 4C).

Neither the application of MMP2 nor MMP13 (1 nM) evoked a rise in intracellular [Ca^2+^] in sensory neurons (p = 0.18 and p > 0.99, respectively, compared to vehicle, Figure 6). Given MMP2 and MMP13 have been shown to cleave PAR1, we wondered if MMP2 and MMP13 could prevent PAR1 activation by other MMPs or TRAP6. To test this, neurons were treated with MMP2 for 10 minutes prior to the application of either MMP3 or TRAP6. If MMP2 does cleave PAR1, then it would be expected that the responses elicited by MMP3 or TRAP6 would be reduced as PAR1 is non-functional (and is internalised) following cleavage. The fraction of MMP3-responsive neurons was reduced by pre-treatment with MMP2 (F(5, 24) = 48.54, p < 0.0001), as was the fraction of TRAP6-responsive neurons (p < 0.0001, Figure 6A). That said, the fraction of neurons stimulated by MMP3 and TRAP6 following MMP2 pre-treatment remained greater than that stimulated by vehicle alone (p = 0.0009 and p = 0.0008, respectively, Figure 6A). As such, a more prolonged pre-treatment (30 minutes) with MMP2 was tested. Again, pre-treatment with MMP2 reduced the fraction of neurons responding to application of MMP3 (F(5, 24) = 63.34, p < 0.0001) and TRAP6 (p < 0.0001, Figure 6B). In this case, the fraction of neurons responding to MMP3 and TRAP6 following treatment with MMP2 was no different to vehicle (p > 0.99 and p = 0.82, respectively, Figure 6B). We performed the same experiments with MMP13 and found very similar results. Pre-treatment (30 minutes) with MMP13 inhibited the subsequent response to MMP3 (F(5, 24) = 12.44, p = 0.0055) and TRAP6 (p = 0.0005, Figure 6C).

**Figure 6:**
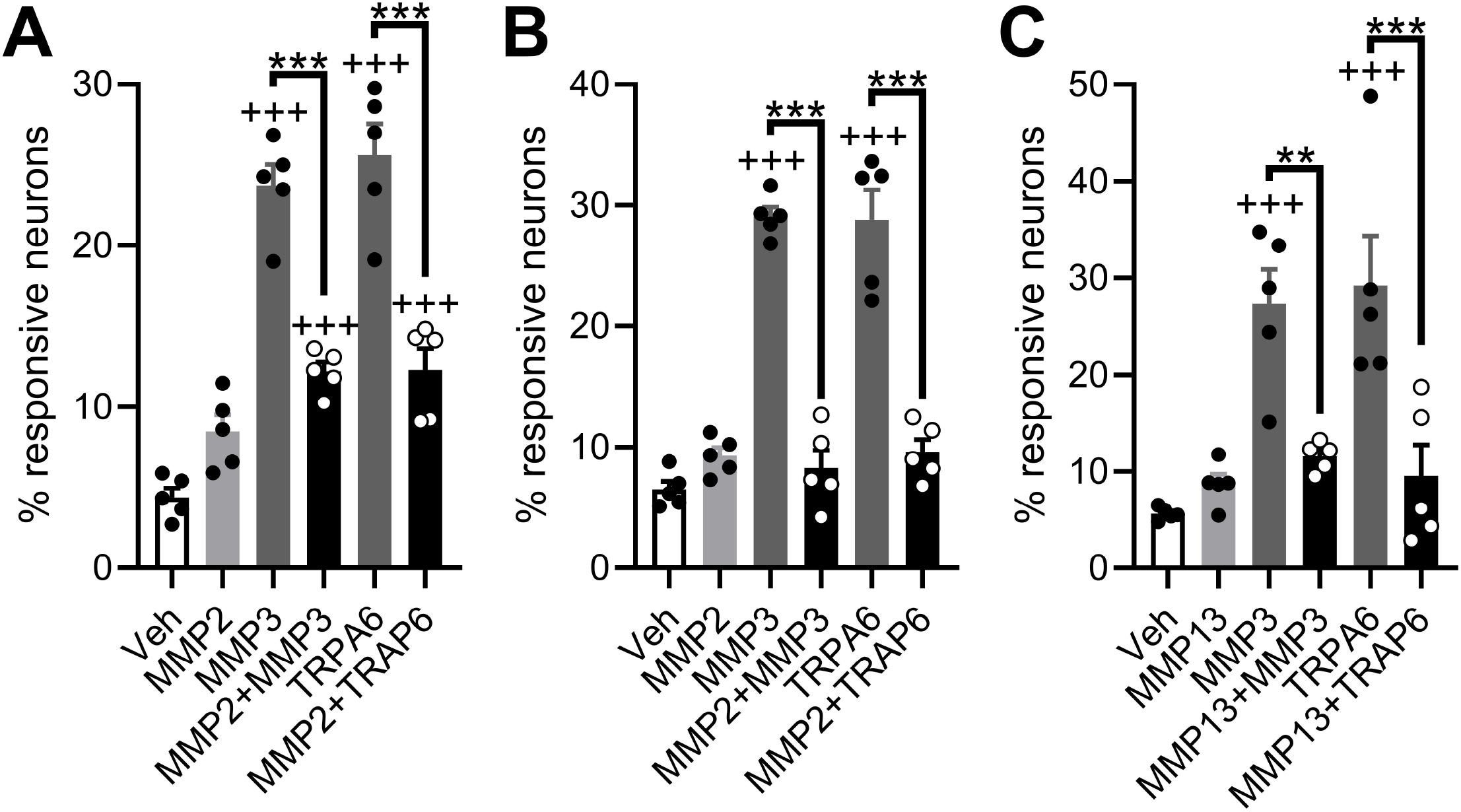
MMP2 and 13 cleave PAR1 but do not evoke a rise in cytosolic Ca^2+^ concentration. (A) Grouped data showing the proportion of neurons stimulated by vehicle, MMP2 alone, MMP3 alone, MMP3 after 10 minute treatment with MMP2, TRAP6 alone, and TRAP6 after 10 minute treatment with MMP2 (5 independent experiments per group). One-way ANOVA with Bonferroni’s *post-hoc* tests. *** show p-values for highlighted comparisons; +++ show p-values for comparisons to the vehicle group. (B) As in (A), except MMP2 pre-treatment was 30 minutes (5 independent experiments per group). One-way ANOVA with Bonferroni’s *post-hoc* tests. (C) Grouped data showing the proportion of neurons stimulated by vehicle, MMP13 alone, MMP3 alone, MMP3 after 10 minute treatment with MMP13, TRAP6 alone, and TRAP6 after 10 minute treatment with MMP13 (5 independent experiments per group). One-way ANOVA with Bonferroni’s *post-hoc* tests.

## Discussion

In this study, multiple members of the MMP family were found to be upregulated in colonic biopsies from patients with Crohn’s disease (Higham et al., 2024). Ca^2+^ imaging revealed that MMP3, 8 and 9 activated PAR1 which induced PLC-dependent intracellular Ca^2+^ release in a proportion of likely nociceptive neurons. In contrast, MMP2 and 13 did not induce Ca^2+^ mobilisation in DRG neurons. However, further experimentation suggested that MMP2 and 13 likely cleave PAR1 as pre-treatment of neurons with MMP2 or 13 reduced neuronal responses to subsequent PAR1 agonist exposure. Additionally, MMP10 induced Ca^2+^ mobilisation in DRG neurons but through a PAR1-independent mechanism. Collectively, these findings provide new insights into the interactions between MMPs and sensory neurons and identify a novel pro-nociceptive signalling pathway in the inflamed bowel.

Our data indicate that the neurons sensitive to MMP3, 8 and 9 are likely nociceptors, based on their soma size, sensitivity to capsaicin and staining with IB4. The majority of MMP-sensitive neurons exhibited a soma area less than 1000 μm^2^, with a median between 400-600 μm^2^. Sensory neurons with soma areas within this range are very likely to be Aδ- and C-type nociceptors (Lawson et al., 2019). Beyond this, sensitivity to capsaicin, an agonist of the heat- and acid-sensitive cation channel, TRPV1, is a key marker of a subset of nociceptive sensory neurons (Dubin & Patapoutian, 2010). Around one-third of the neurons which responded to the application of MMP3, 8 or 9 also responded to capsaicin. This provides further evidence that MMP-sensitive neurons are nociceptors. Finally, IB4 staining revealed approximately one-half of MMP-sensitive neurons were positively stained with IB4, indicating that they are likely non-peptidergic nociceptors (Fullmer et al., 2004; Pinto et al., 2019). Within this MMP-sesnsitive non-peptidergic population, the majority were insensitive to capsaicin, indicating that these neurons are likely to express *Mrgprd* (Usoskin et al., 2015). It has been demonstrated that *Mrgprd*-expressing sensory neurons contribute to nociceptive signalling in the colon (Bautzova et al., 2018). These data strongly suggest that MMP3, 8 and 9 evoked a rise in cytosolic Ca^2+^ concentration in a subset of nociceptive sensory neurons *in vitro*.

Previous reports have indicated that multiple members of the MMP family, including MMP1, 3, 8 and 9, cleave PAR1 (Boire et al., 2005; Lee et al., 2010; Trivedi et al., 2009). To test the involvement of PAR1 in Ca^2+^ signals evoked by MMP3, 8 and 9 in sensory neurons, we used the PAR1-selective antagonist, vorapaxar (Hawes et al., 2015). We verified that vorapaxar inhibits PAR1 by demonstrating that the neuronal response to TRAP6, a PAR1-selective agonist, was inhibited by pretreatment with vorapaxar. The inhibitory effect of vorapaxar in our experiments indicates that PAR1 is required for MMP3-, 8- and 9-evoked Ca^2+^ signals in sensory neurons. We also found that another PAR1 antagonist, SCH79797, inhibited MMP1-evoked Ca^2+^ signals, suggesting that PAR1 may be a common receptor for MMPs. Downstream of PAR1 activation, our data indicates that Ca^2+^ signals rely on PLC-mediated Ca^2+^ release from intracellular stores. This is in line with previous data from sensory neurons showing that PAR1-mediated Ca^2+^ signals are unaffected by the removal of extracellular Ca^2+^ (Amadesi et al., 2004), and data from fibroblasts demonstrating a requirement for PLC and replete intracellular Ca^2+^ stores for thrombin-evoked Ca^2+^ signals (Jeng et al., 2004). Biopsy supernatants derived from patients with irritable bowel syndrome stimulated sensory neurons in a PAR1-dependent manner, implicating PAR1 in pro-nociceptive signalling (Desormeaux et al., 2018).

We observed no effect of PAR1 stimulation – either with MMP3, 8, 9 or TRAP6 – on the neuronal response to capsaicin (data not shown). There is conflicting evidence concerning the sensitisation of TRPV1 by PARs in sensory neurons. For example, it has been reported that both PAR1 and PAR4 sensitised TRPV1 (Vellani et al., 2010), but another report showed that PAR2, not PAR1, sensitised TRPV1 (Amadesi et al., 2004). Our data aligns with the latter example, though the reasons underlying these conflicting results are not immediately clear.

While MMP10 did evoke Ca^2+^ signals in sensory neurons, we observed no effect of vorapaxar on the response to MMP10. To the best of our knowledge, MMP10 has not previously been shown to cleave PAR1 (Heuberger & Schuepbach, 2019). Perhaps MMP10 signals through another member of the PAR family. Agonists of PAR2 and PAR4 evoke Ca^2+^ signals in sensory neurons (Vellani et al., 2010), so future work could examine the involvement of these receptors in the response to MMP10.

In contrast to MMP1, 3, 8, 9 and 10, MMP2 and 13 did not evoke Ca^2+^ signals in sensory neurons. MMP2 has been shown to induce slowly-developing Ca^2+^ release in platelets in a PAR1-dependent manner. What’s more, MMP2-mediated cleavage of PAR1 is more efficient in the presence of the α_IIb_β_3_ integrin (Sebastiano et al., 2017), which is poorly expressed in sensory neurons (Zeisel et al., 2018). As such, our assay may have been of insufficient duration to capture MMP2-evoked Ca^2+^ signals. It is also possible that MMP2-mediated cleavage reveals a biased tethered ligand which does not stimulate Ca^2+^ mobilisation. That said, we have still garnered evidence that MMP2 cleaves PAR1 expressed by sensory neurons, because pretreatment with MMP2 inhibited the subsequent response to either MMP3 or TRAP6. Similarly, we did not observe Ca^2+^ signals evoked by MMP13. A previous report showed that, in cardiac fibroblasts, MMP13 induced PAR1-dependent ERK1/2 activation, as well as PAR1 internalisation, but did not evoke IP_3_ production, indicating that MMP13 may not stimulate Ca^2+^ mobilisation (Jaffré et al., 2012). The inhibition of the neuronal response to MMP3 or TRAP6 by pretreatment with MMP13 provides further evidence that MMP13 does cleave PAR1.

Given the risk of bleeding, PAR1 antagonists are unlikely to be of use in the treatment of visceral pain. Broad-spectrum MMP inhibitors are associated with severe side effects, such as musculoskeletal syndrome (Fingleton, 2008), but more selective inhibitors, like tanomastat, are not (Das et al., 2021). As such, selective MMP inhibitors may have potential as treatments for visceral pain in IBD. That said, our study can only provide information on the effects of MMPs in isolation. The environment in the inflamed bowel is highly complex with myriad interacting processes contributing to nociception and pain (Roda et al., 2020). To understand the potential for MMP inhibitors as treatments for visceral pain, it will be necessary to first understand the role of MMPs in the inflamed bowel. Proteolytic activity, including that of thrombin, is increased in the inflamed bowel (Motta et al., 2021; Steck et al., 2012), and the expression of various MMPs is correlated with the extent of inflammation (Von Lampe et al., 2000) – indicating a potential role for MMPs in IBD pathophysiology. MMPs are upregulated in other inflammatory conditions, such as rheumatoid arthritis and psoriasis (Mezentsev et al., 2014; Pulik et al., 2023), and so may contribute to pro-nociceptive signalling in these conditions too.

This study has highlighted a potential pro-nociceptive role for multiple members of the MMP family which are elevated in the inflamed bowel in Crohn’s disease. This broadens our understanding of the mediators and mechanisms underpinning nociception and pain during colonic inflammation.

